# Integrated Disease Surveillance and Response (IDSR) in Malawi: Implementation Gaps and Challenges for Timely Alert

**DOI:** 10.1101/363713

**Authors:** Tsung-Shu Joseph Wu, Matthew Kagoli, Jens Johan Kaasbøll, Gunnar Aksel Bjune

**Affiliations:** Department of Informatics, University of Oslo, Norway; Research Department, Luke International, Malawi; Overseas Mission Department, Pingtung Christian Hospital, Taiwan; Department of Public Health, National Taiwan University, Taiwan; Department of Epidemiology, Ministry of Health, Malawi; Public Health Institute, of Malawi, Malawi; Institute of Health and Society, University of Oslo, Norway

## Abstract

**Objective:** The emerging and recent 2014 Ebola Virus Disease (EVD) outbreaks rang the bell to call upon efforts from globe to assist resource-constrained countries to strengthen public health surveillance system for early response. Malawi adopted the Integrated Disease Surveillance and Response (IDSR) strategy to develop its national surveillance system since 2002 and revised its guideline to fulfill the International Health Regulation (IHR) requirements in 2014. This study aimed to understand the state of IDSR implementation and differences between guideline and practice for future disease surveillance system strengthening.

**Methods:** This was a mixed-method observational study. Quantitative data were to analyze completeness and timeliness of surveillance system performance from national District Health Information System 2 (DHIS2). Qualitative data were collected through interviews with 29 frontline health service providers from the selected district and key informants of the IDSR system implementation and administration at district and national levels.

**Findings:** The current IDSR system showed relatively good completeness (76.4%) but poor timeliness (41.5%) of total expected monthly reports nationwide and zero weekly reports. The challenges of IDSR implementation revealed through qualitative data included lack of supervision, inadequate resources for training and difficulty to implement weekly report due to overwhelming paperwork at frontline health services.

**Conclusions:** The differences between IDSR technical guideline and actual practice were huge. The developing information technology infrastructure in Malawi and emerging mobile health (mHealth) technology can be opportunities for the country to overcome these challenges and improve surveillance system to have better timeliness for the outbreaks and unusual events detection.

## Introduction

After the largest Ebola Viral Disease (EVD) outbreak happened in Western Africa, governments, health authorities in Africa and the world learnt a valuable lesson from the challenges of diseases surveillance systems implementations in countries with limited public health infrastructure [1, 2]. The outbreak emerged in 2013, ended in June 2016 and affected 10 countries worldwide with 28,616 confirmed or probable cases, and 11,310 deaths [3-6]. The feebleness of the public health infrastructure and capabilities, to capture early warning signal of outbreak and provide good timeliness for response, was further exposed during this epidemic and the need for strengthening the surveillance system in these countries and transform it from passive to active surveillance was articulated for actions [2, 7]. Yet the new EVD outbreak emerged in the Democratic Republic of the Congo in April 2018 [8].

Early case detection is one of the important approaches to managing future outbreaks [9]. In Africa, although Integrated Diseases Surveillance and Response (IDSR) strategy was adopted as the regional development approach for member states and technical partners since 1998, still, challenges of implementing IDSR have highlighted already before the tragic EVD outbreak event in 2014 [10-12]. Following the Severe Acute Respiratory Syndrome (SARS) outbreak in 2003, the International Health Regulation (IHR) was revised by the World Health Organization (WHO) in 2005 and fully adopted by all countries around the world [13]. The IHR-2005 enhancement proved to be helpful in dealing with the 2009 H1N1influenza pandemic and IDSR serves the platform for its implementation in Africa [14, 15]. However, shortcomings of the global health system’s capability, lack of virological surveillance in Africa and technologies for vaccine production and implementation and the basic public health system infrastructure were revealed during the same pandemic [16].

Malawi adopted the IDSR in 2002 and the third edition technical guideline was published in May 2014 with incremental reportable diseases and health conditions to fulfill the IHR-2005 and public health needs [17]. The epidemiology department (ED) of the Ministry of Health (MOH) is the main custodian of the IDSR system while the Center for Central Monitoring and Evaluation Division (CMED) in the Department of Planning and Policy Development in the MOH is responsible for coordinating the routine Health Management Information System (HMIS) and its subsystems, including IDSR [18]. The IDSR system reporting and information flow follows the health system organization structure from the community to the national level (Fig 1. The IDSR system information flow according to the organization architecture in Malawi.).

**Figure 1.**
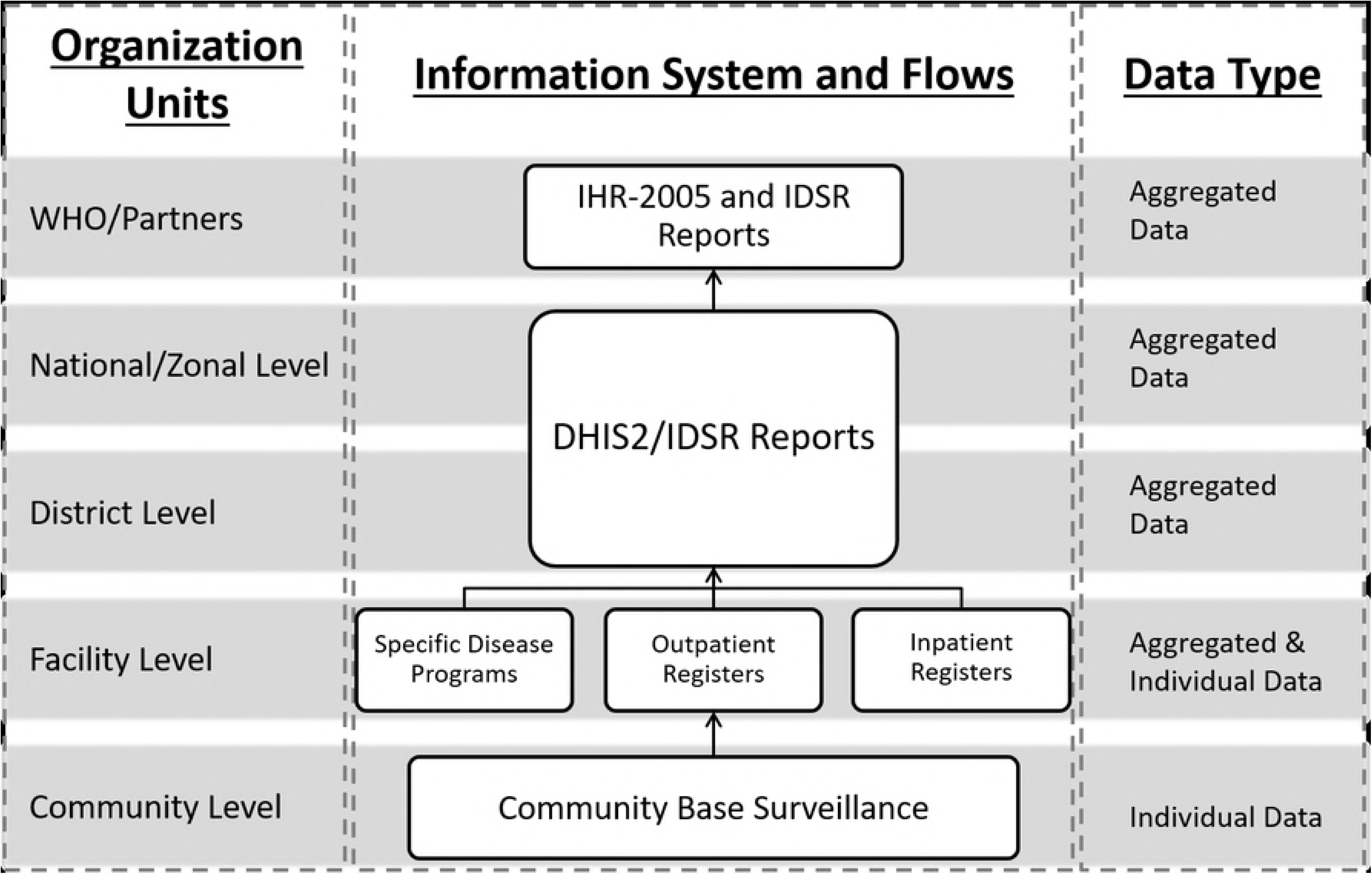
The IDSR system information flow according to the organization architecture in Malawi.

At the community level, the Health Surveillance Assistants (HSAs) are the frontline health care workers (HCWs) responsible for case identification and report. They work under the supervision of attached health facilities to identify case and further refer to the nearest health facility [19]. Most of the health facilities are public, government-owned or non-governmental, non-profit organization under government regulatory in Malawi. The HCWs at each facility, irrespective of ownership (public or private) are responsible for case identification and reporting (weekly and monthly). Each facility has a person responsible for tallying reportable cases using various health information tools, including electronic medical records (EMR) system. In the current guideline, 19 diseases and conditions are required immediately reporting (Table 1).

**Table 1:** Diseases, conditions or events requiring immediate reporting of Malawi IDSR system

| **Table 1: Diseases, conditions or events requiring immediate reporting of Malawi IDSR system [17]** |  |
| --- | --- |
| • Acute Flaccid Paralysis (AFP)<br>• Acute hemorrhagic fever syndrome (Ebola, Marburg, Lassa Fever, Rift Valley Fever (RVF), Crimean-Congo)<br>• Adverse effects following immunization (AEFI)<br>• Anthrax<br>• Cholera<br>• Cluster of SARI<br>• Diarrhoea with blood (Shigella dysentery)<br>• Influenza due to new subtype<br>• Maternal death<br>• Measles | • Meningococcal meningitis<br>• Neonatal tetanus<br>• Plague<br>• Rabies (confirmed cases)<br>• Severe Acute Respiratory Syndrome (SARS)<br>• Smallpox<br>• Typhoid fever<br>• Yellow fever<br>• Any public health event of international concern (infectious, zoonotic, food borne, chemical, radio nuclear or due to an unknown condition) |

Each District Health Office (DHO) has a District Health Management Team (DHMT) overseeing health programmes. The District Environmental Health Officer (DEHO) of DHMT is responsible for HSAs management and district IDSR focal person is collecting surveillance reports from facilities for submission and notification. From district level above, Malawi has adopted District Health Information System (DHIS) as the national system for HMIS reporting since 2002. The system was upgraded to a web-based open-source information system, DHIS2, in 2012 [20]. MOH is hosting DHIS2 and the IDSR reports are required to be entered by the focal person since late 2014. The IDSR core functions of each levels of the health system clearly articulated in the guideline (Table 2).

**Table 2.** The Malawi IDSR core functions and activities at each health system level

| Level of Health System | Core Functions |  |  |  |
| --- | --- | --- | --- | --- |
|  | Identify | Report | Analysis and Interpret | Investigate and Confirm |
| **Community** | Use simple case definitions to identify priority diseases, public health events, conditions or other hazards in the community | Report case critical information on priority diseases, public health events, conditions, or hazards to health facility and appropriate authorities | Involve local leaders in observing describing and interpreting disease patterns, public health events and trends in the community.<br>Undertake verbal autopsies on causes of deaths | Support event investigation activities. |
| **Health Facility** | Use standard case definitions to detect confirm and record priority diseases or conditions public health events<br>Collect and transport specimens for laboratory confirmation<br>Use local laboratory capacity to confirm cases or to initiate confirmation of cases if possible | Report case based information for immediately notable diseases<br>Report summary data to next level<br>Report laboratory results from screening of sentinel populations<br>Report laboratory results to next level | Prepare and periodically update graphs tables, and charts to describe time, person and place for reported diseases, conditions and public health events<br>From the analysis, report immediately any disease or condition that:<br>exceeds an action threshold<br>occurs in locations where it was previously absent<br>presents unusual trends or patterns<br>Interpret results, initiate possible Public Health actions with local authorities | Take part in investigation of reported outbreaks or public health events<br>Collect, package, store and transport specimens for laboratory confirmation |
| **District, Zonal** | Collect surveillance data from reporting sites including designated points of entry on time and review the quality<br>Ensure reliable supply of data collection and reporting tools are available at reporting sites<br>Collect and transport specimen for laboratory confirmation<br>Use local laboratory capacity to confirm cases if possible | Make sure health facilities know and use standard case definitions for reporting priority diseases, conditions and public health events<br>Make sure health facility staff know when and how to report priority diseases, conditions and public health events<br>Report data on time to the next level<br>Report laboratory results to next level<br>Periodically conduct risk assessment for priority diseases, | Define denominations and ensure their accuracy<br>Aggregate data form health facility reports<br>Analyze data by time, place and person<br>Periodically update graphs, tables and charts to describe reported diseases, public health events and conditions<br>Calculate rates and thresholds<br>Compare current data with previous periods<br>Make conclusions about trends, | Arrange and lead investigation of reported diseases or outbreaks and public health events<br>Assist health facility in safe collection, packaging storage and transport of laboratory specimen for confirmatory testing<br>Receive and interpret laboratory results<br>Describe if the reported outbreak and public health events is confirmed.<br>Report the confirmed outbreak or public health events to the next |
|  |  | public health events, condition or hazards | thresholds and analysis results<br>Describe risk factors for priority diseases or conditions or public health events | level<br>Distribute specimen collection Kits for special surveillance activities. |
| **NATIONAL** | Define, update and ensure compliance with national policy and guidelines<br>Set policies and procedures for the national laboratory networks including quality assurance systems<br>Use national laboratory for confirmatory and specialized testing if necessary<br>Collect and transport specimens for additional analysis at WHO Collaborating Centres as necessary | Report the immediate notifiable diseases and public health events to the appropriate authorities on time<br>Report other priority disease and public health events on time<br>Include all relevant laboratories in the reporting network<br>Use IHR Decision Instrument (Annex2) to determine risks for priority diseases public health events, conditions or hazards<br>Inform WHO as indicated by IHR (2005) | Set policies and procedures for analyzing and interpreting data<br>Analyze and interpret data from a national perspective<br>Regularly convene a meeting of the technical coordinating committee to review the analyzed and interpreted data before wider dissemination<br>Carry out special analyse to forecast magnitude and trends of priority disease and public health events. | Ensure guidelines and standard operating procedures for outbreak investigations are available at all sites<br>Coordinate and collaborate with international authorities as needed during investigations<br>Alert and support laboratory participation<br>Provide logistic support l supplies, equipment, reagents, specimen transport media, health promotion budget<br>Share information with regional and international networks about confirmed outbreak<br>Use the decision instrument to decide whether the outbreak is a potential PHEIC<br>Process specimen from investigation and send timely results as required to each level<br>Request additional specimen as needed<br>Take part in epidemic response team |

Despite existing framework of IDSR system, few nationwide assessments of IDSR system have been done in Africa and none in Malawi [12, 21-23]. This study aims to explore the differences between the IDSR guideline and practice, specifically looking into the timeliness and completeness aspects, which shall trigger responses.

## Materials and Methods

### Study design

This study mixed quantitative and qualitative methods to assess and understand the implementation gaps of IDSR system from each level of the health system in Malawi and focused on two key attributes, timeliness and completeness, of the surveillance system [24].

### Source of data

We used the built-in function of the DHIS2 to extract IDSR monthly reporting rate summary data from the central server of the Ministry of Health, period from October 2014 to September 2016 and the data were extracted in June 2017.

Qualitative data of community to district level IDSR workers were collected from one convenience selected district in the Northern Region of Malawi, which has the best performance of IDSR reporting in 2013. The interviews and observations were conducted based on the interview guide and conducted in English, Chewa or Tumbuka. The interviews were digitally recorded for transcribing, translating and analysis. The researcher (TSJW) observed operation of the outpatient clinic in hospitals to obtain contextual information about the service providing and IDSR report generating process. Key informants from the district and the national level were interviewed to obtain IDSR system implementation status and to identify the challenges and gaps.

### Data analysis

Quantitative data were exported from the DHIS2 with the Excel data format and divided by the studied district and one national category. The data were compiled to one dataset and analyzed using tabulation and line charts to illustrate the time series patterns of the IDSR monthly report data quality – completeness and timeliness^1^. According to the national policy, 80% completeness and timeliness is the threshold of good performance.

Interview records were transcribed into text for translating and read by the researchers (TSJW) to find the actual practices of IDSR system. The core functions of each level of health system actors were compared with the expected functions according to the guideline.

## Results

### Quantitative data

We extracted 168 IDSR reporting rate summary (24 months). The completeness data, exclude the outlier in February 2015, showed average completeness was 94.0% and 76.4% in the studied district and nationwide respectively (Fig 2. The IDSR monthly reports completeness indicator from October 2014 to September 2016 divided by national level and studied district). Only 4.2% of the IDSR monthly reports from the whole country reached the good performance standard.

**Figure 2.**
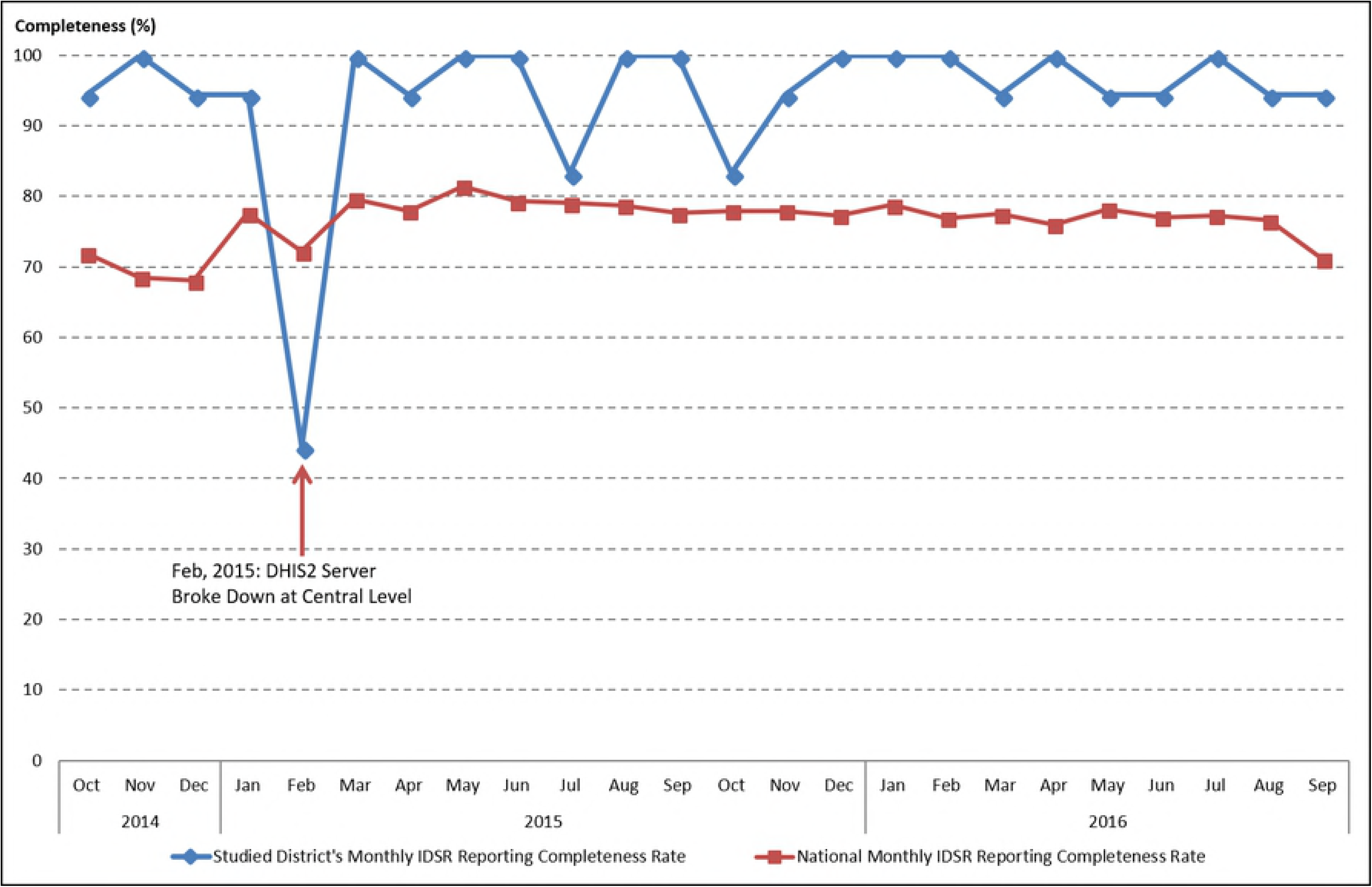
The IDSR monthly reports completeness indicator from October 2014 to September 2016 divided by national level and studied district.

We observed very poor timeliness performance of the IDSR monthly report (Fig 3. The IDSR monthly reports timeliness indicator from October 2014 to September 2016 divided by national level and studied district). Good performance was not achieved during any of the 24 months.

**Figure 3.**
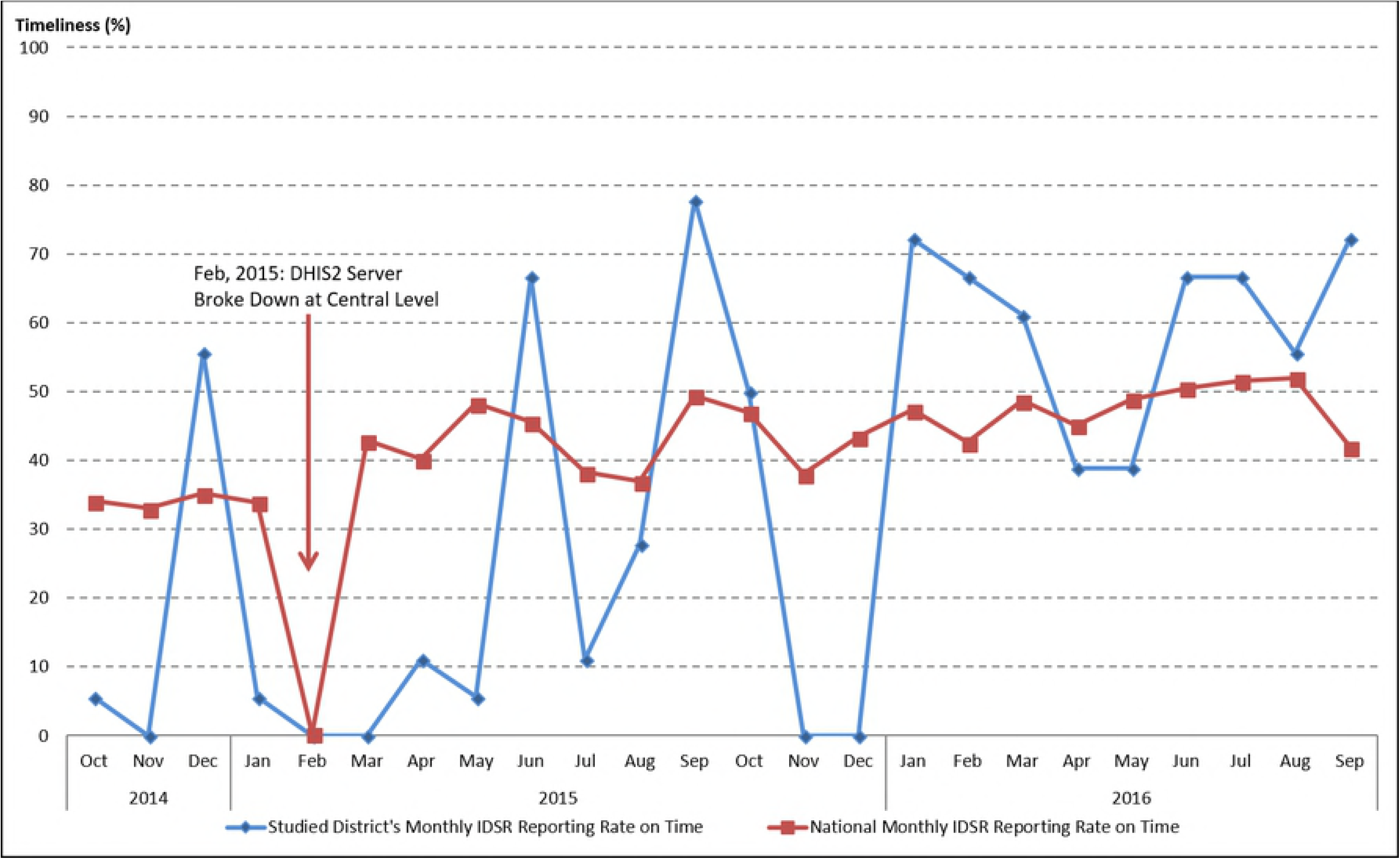
The IDSR monthly reports timeliness indicator from October 2014 to September 2016 divided by national level and studied district.

Notably in February 2015, the timeliness of IDSR monthly reports all dropped to almost 0% due to server breakdown and also affected the completeness of IDSR monthly report in the studied district.

### Qualitative data

#### Community level health workers

At the community level, we interviewed 17 HSAs who run the village clinics to provide health services to the villagers. However, according to all the informants, none of them was practicing community-level case identification using IDSR guidelines. They relied on volunteers from the Village Health Committee (VHC) to report unusual health events. One informant explained the limited logistic support and large catchment area to serve constitute challenges to do active surveillance works. The health volunteers from the VHC hence played critical roles for the community level outbreak or incidence alerts.

> *“I have volunteers from each village, 2 of them* (in each village). *Those volunteers are my ambassadors. They have the knowledge, if any outbreak, they tip me, then I rush* (to the village).*”* HSA, #RU03

They initiated preliminary investigations when community rumors emerge and physically walked to the higher-level health facilities to report.

> *“I can write a written report then submit it to office, or I can go in person explain the situation to my boss.”* HSA, #RU02

The main function that HSAs saw for themselves in serving IDSR was to assist their health facilities to compile the IDSR monthly reports, perform community sensitization and education. Through observations, some HSAs were equipped and capable to do simple data analysis (Fig 4. Population statics tabulated by health surveillance assistant in one village clinic) and use them as instrument to interact with the VHC for disease prevention and health promotion. However, the limited supervision and resources affected their performance in this area of work.

> *“At first, we are going to each and every household…but this time it’s not often.”* HSA, Informant #RU03

**Figure 4.**
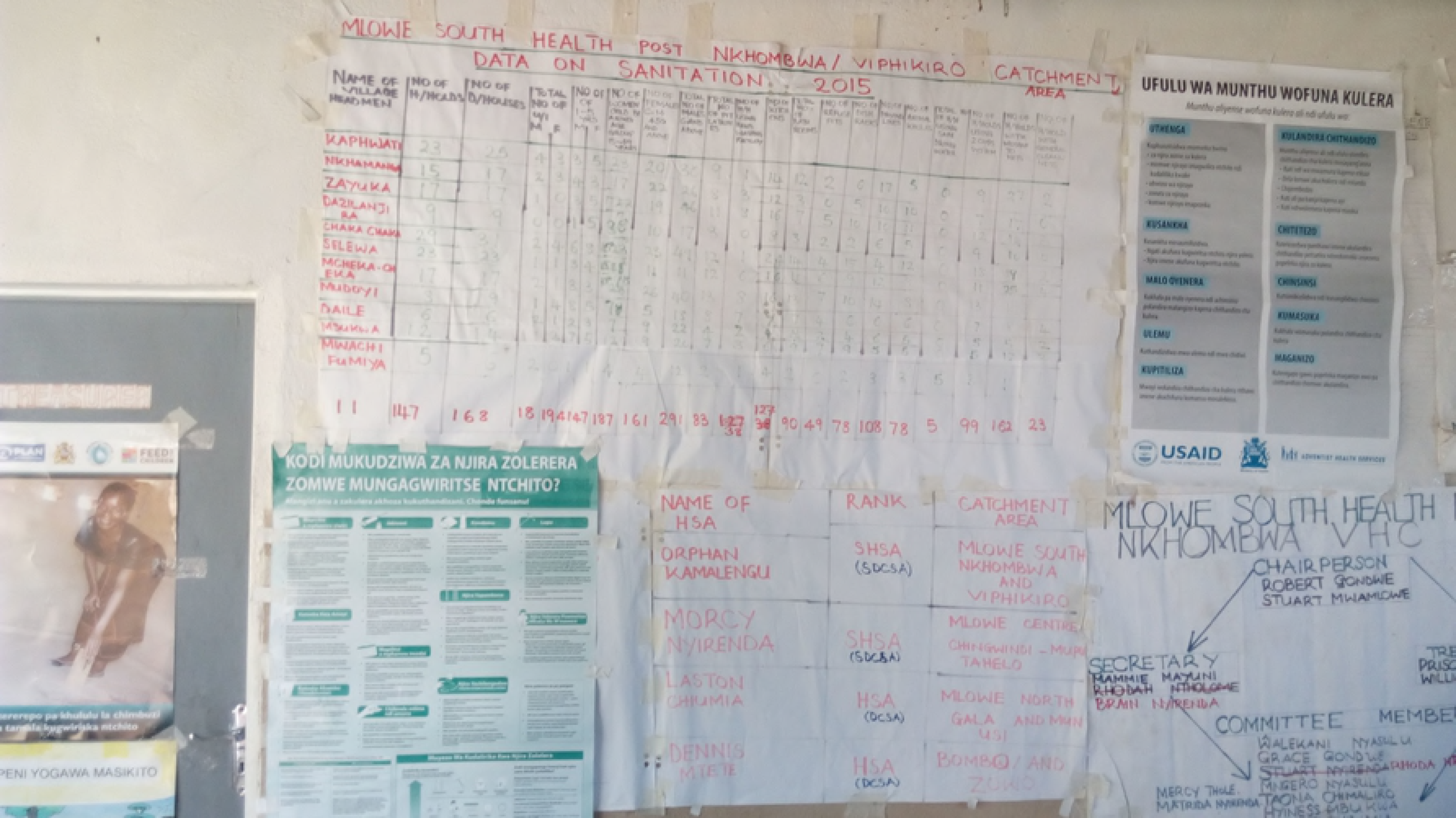
Population statics tabulated by health surveillance assistant in one village clinic

#### Facility level health care workers

At the facility level, we interviewed 12 HCWs from two health facilities. The HCWs picked up unusual health events through their daily services and did not wait until the monthly report to take actions.

> *“When we see many patients are coming from that place and they are registering ARI* (acute respiratory infection). *What does that mean? So we don’t even wait for the month to come and work on the data. But we just see that I think for this*… *we come together and then we discuss. If it’s an outbreak we see that we cannot control, then we inform the DHO.”* HCW, #MHRH01
>
> *“We usually report to the environmental officer and then they will send the HSAs and see what’s going on there.”* HCW, #RDH04

Despite of EMR system in place, the heavy workload made it difficult for HCWs to capture clinical information on system for automate reporting. They simplified the work and transcribed individual level data into the different paper registers for reporting.

> *“The computers are not fast as we expected them to be. Just to print somebody’s name, you have to wait for a minute or more. So you say this is delaying me, let me just write*… *we are having a lot of patients. A lot of them.”* HCW, #RDH04

#### District level

The challenges to get timely reports through unstable information technology infrastructure were obstacles for the IDSR focal person in the DHO to provide quality reports.

> *“*…*with IDSR, I have got challenges with the reporting system itself, from the health facilities, sometimes reports come a bit late. We also have challenges of that we do not have internet at the hospital. So we have to use the smart phones, the* (internet) *dongles to buy units and we are not provided with any funding for internet services so we have got to go into our pockets*…*”* DHO, #RDHO02

Lack of comprehensive training was the challenge to enhance the electronic system to capture more data for disease surveillance and decision-making.

> *“The challenge is those who are using the computers, it’s just a few number of people who are oriented*… *that’s why it’s difficult to capture the data and many information.”* DHO, #RDHO02

Financial constrains were key concerns. This created gaps for IDSR system to be implemented using the updated technical guideline at the community and facility levels.

> *“The new IDSR guidelines are in, but due to lack of funds they have not yet called us for orientation on the new guidelines. We are still using the old guidelines, which is having fewer information.* … *We are just waiting* (fund) *so that we can also share the information with our fellow health workers.”* DHO, #RDHO02

#### National level

We noted during the field study at community, facility and district level that no one mentioned the IDSR weekly reports, nor actually implementation of the new guideline. The constrained resources heavily affect the performance of the IDSR system in the country.

> *“At the beginning we are doing very well. WHO came and helped us to setup the system from 2002, we do supervision, training and so on, up until 2007 there is no fund. Government said we cannot take it, it’s too costly.”* ED, #MOH02

The IDSR weekly reporting system paralyzed due to the difficulties for HCWs to cope with the volume of paper-generated reports and lack of internet connectivity. This seemed the main obstacle from national authorities perspective who eagers to enable the system for rapid responses.

> *“Of course we told them to do weekly report, but there is no internet. For them to write report and send*… *it is just too difficult for them to handle these papers.”* ED, #MOH02

The data quality of IDSR monthly report submitted through HMIS was the concern for ED to use. For instance, there were 31 Viral Hemorrhagic Fever cases recorded in DHIS2 in 2015, but none confirmed by the department.

> *“If you look at the data, you will be surprised like: how can we have Ebola cases and we don’t know. The data quality is just so poor and we cannot use it.”* ED, #MOH01

The department expected to use technology to improve timeliness and capability to early response.

> *“If it is an immediately notifiable case, we want to know immediately. We don’t even want to wait them to report to us, we want to know now. Even it is a rumor or what, we need to know so we can check if it is true. That’s why we want to use this SMS or the eIDSR* (electronic IDSR) *so we can know there is something happening there.”* ED, #MOH02

There is a fundamental difference between the needs of the HMIS and the IDSR systems, where one is looking only for confirmed cases while IDSR is looking for alerts to take fast actions.

> *“We want to get confirmed cases. We need to know exactly how many are they so we can do proper planning. That is why we want the data to be complete and accurate.”* CMED #MOH03
>
> *“We need to know if there is something happen in the community. Wait for a month, sometimes three months to get report, it is just too slow. We need to take actions immediately so we are looking for any signal that can trigger us to take actions.”* ED, #MOH01

## Discussion

We assessed the differences between IDSR technical guideline and actual practice in the health system in Malawi for the first time. According to the quantitative data, we observed relatively good completeness of IDSR monthly reports compared to timeliness. Timeliness is a general problem to countries implementing IDSR system across Africa, and this makes the public health authorities unable to take quick action and respond to the suspected health events [12, 25]. Facility level IDSR reports may not be sufficiently timely to pick up the outbreaks from community. The strengthened community level surveillance and verbal autopsy to detect unusual deaths can be a good approach to detect lower level health events and provide timely response [26]. In Malawi, a pilot study conducted in Lilongwe District in Central Region showed that mobile technologies had good opportunities to improve timeliness of HMIS reports [27]. However, concerning the different purposes of HMIS and IDSR system, a more integrated electronic IDSR system is essential for the health authorities to correspond the diverse demands.

African health ministries are quickly adopting mHealth solutions to improve disease surveillance and health programmes. Tanzania piloted an IDSR reporting system using SMS function and regular phones for report in 2011 [28] and further expanded it to be the national strategy for diseases surveillance using Unstructured Supplementary Service Data (USSD) technology linked with DHIS2 for the immediate reporting for IDSR [29-31]. Zambia tried to use DHIS2 mobile to enhance its malaria surveillance in Lusaka district and to improve case management and reporting [32]. Other mobile technologies including smartphone applications, patient monitoring devices, Personal Digital Assistants (PDAs), as well as laptops and tablets PCs connected with network service were piloted and implemented in various African countries [33]. Countries and development partners are eager to apply the mobile technology to capture real-time field data for surveillance and case management at the community level health care system [30, 34-37]. However, notable issues were documented including technical, financial, infrastructural challenges, data security and medical supports during the design and implementation process of mHealth surveillance in sub-Saharan Africa countries [33]. Considering the complexity of public health works and needs of integration services at the community level [38], the utilization of mobile technologies requires more rigorous studies to evaluate such innovations for programme implementation to become sustainable and scalable [39].

Apart from mHealth solutions, researchers recommended to use syndromic surveillance approach combined with systematic virological testing as early as possible to maintain high quality situational awareness [40]. Several countries have established electronic data based syndromic surveillance systems to capture early warning signals of different diseases and health status especially related to respiratory infections [41-44]. However, electronic syndromic surveillance systems remain a novel technology for most of developing countries to adopt and implement [45]. Several EMR systems had been developed in Malawi and MOH decided to move towards a national standardized EMR system to support all levels of HMIS [46-48]. This provides a unique opportunity to utilize existing information technology and infrastructures to strengthen the IDSR system with nationwide syndromic surveillance. Yet it is critical to improve the user experiences of EMR users to improve the uptake and usage of the system. Similar countries can consider system synergies and existing infrastructure for IDSR enhancement.

We only focused on completeness and timeliness, and the accuracy attribute of the IDSR system performance was out of the scope of this study. Further clinical and laboratory data are needed for proper assessment. We only sampled one district to conduct qualitative assessment, however, we are confident that it is relevant for the Malawian context by the fact that the health care system is rather homogeneous in Malawi and the district we selected had a relatively good IDSR performance to generalize the implementation challenges.

## Conclusions

Lack of timeliness in reporting makes the IDSR system inoperative. Differences between IDSR technical guideline and actual practice existed in the current Malawian context. Shortcomings were due to financial constraints and poor basic infrastructure. However, the improving information technology infrastructure in Malawi, single country platform EMR system and emerging mHealth technologies can be opportunities for the country to overcome the challenges and improve the surveillance system.

## Acknowledgements

Omega Banda the research assistant who assisted the filed study data collection, transcribing and translation of the materials.

## List of abbreviations

AFRO: World Health Organization Regional Office for Africa,
ARI: acute respiratory infections,
CDC: Centers for Disease Control and Prevention,
CHAM: Christian Health Association of Malawi,
CMED: central monitoring and evaluation division,
DEHO: district environmental health officer,
DHIS: district health information system,
DHIS2: district health information system 2,
DHMT: district health management team,
DHO: district health office,
EMR: electronic medical records,
EVD: Ebola virus disease,
HCWs: health care workers,
HMIS: health management information system,
HSAs: health surveillance assistants,
IDSR: integrated disease surveillance and response,
IHR: international health regulation,
mHealth: mobile health,
MOH: Ministry of Health,
PDAs: Personal Digital Assistants,
SARS: severe acute respiratory syndrome,
USSD: unstructured supplementary service data,
VHC: village health committee,
WHO: World Health Organization.

## Declarations

### Ethics approval and consent to participate

The study protocol was reviewed and approved by the National Health Sciences Research Committee (NHSRC) of Malawi with approval number 16/4/1563. The study was granted permission by the health authorities from district health office and the Ministry of Health. All interviews were conducted with the written consent from the interviewees.

### Consent for publication

Not applicable, no individual patient data is included in this study.

### Availability of data and materials

The datasets generated and analyzed during the current study are not publicly available due the DHIS-2 is Malawi government owned internal dataset and personal interviews but are available from the corresponding author on reasonable request.

### Authors’ contributions

TSJW analyzed and interpreted the data. MK contributed IDSR evaluation direction and policy interests from the government prospective. JJK and GAB contributed to the structure and argument directions of the study and analysis. All authors read and approved the final manuscript.

## Footnotes

1 Completeness of reporting indicates whether facilities have reported on the IDSR monthly data they are supposed to report on, while timeliness indicates whether these reports were delivered on time. According to the national policy, each health facility has to compile the IDSR monthly report by 15^th^ of the month and the districts received and entered by 25^th^ of the month.

